# Retention of Lumpy Skin Disease Virus in *Stomoxys* Spp (*Stomoxys Calsitrans, Stomoxys Sitiens, Stomoxys Indica*) following intrathoracic inoculation, Diptera: Muscidae

**DOI:** 10.1101/2020.08.13.249227

**Authors:** Arman Issimov, Lespek Kutumbetov, Assylbek Zhanabayev, Nurlybay Kazhgaliyev, Birzhan Nurgaliyev, Izimgali Zhubantayev, Aliya Akhmetaliyeva, Malik Shalmenov, Abzal Kereyev, Peter J. White

## Abstract

Lumpy skin disease (LSD) is an emerging disease in cattle in Kazakhstan and the means of transmission remains uncertain. In the current study, acquisition of Lumpy Skin Disease Virus (LSDV) by *Stomoxys* species following intrathoracic inoculation was demonstrated under laboratory conditions. Flies were injected with a virulent LSDV strain into the thorax region to bypass the midgut barrier. The fate of pathogen in the hemolymph of the flies was further examined using PCR and Virus isolation tests. LSDV was isolated from all three *Stomoxys* species immediately and up to 24h post intrathoracic inoculation while virus DNA was detectable up to 7d post intrathoracic inoculation.

## Introduction

Lumpy Skin Disease (caused by LSDV, double-stranded DNA virus, family *Poxviridae*) is an economically devastating disease of cattle (1). It is a highly contagious skin disease mainly characterized by multiple skin lesions, fever, enlargement of superficial lymph nodes, excessive salivation, lacrimation and nasal discharge as well as oedema and swelling of the limbs. The World Organization for Animal Health (OIE) classified LSD as a notifiable disease due to its significant economic impact (2). In addition, it has a detrimental effect on animal production. Economic loss results from a sharp decline in milk yield, milk quality, hide damage, body weight reduction, abortion, infertility and in some cases death of the animal (3). Morbidity rates may vary significantly during LSD outbreaks and reach up to 100%, whereas the mortality rate is usually low (less than 5%) reaching 20% on some occasions (2, 4, 5).

Stable flies (*Stomoxys* spp) have long been considered a major pest of livestock in Kazakhstan and are capable of transmitting pathogens present in infected animals. (6). A high concentration of these flies is observed in almost all regions in Kazakhstan with adult *Stomoxys* flies emerging in large numbers following rainfall events in the later months of spring.

To date, LSDV transmission routes and possible insect carriers remain uncertain; however, in controlled studies, mechanical transmission of LSDV was successfully demonstrated using *Stomoxys* species (7, 8). In their study, the presence of LSDV in *Stomoxys* flies was recorded for a short time ranging from 0 – 48 hours post feeding. LSDV was detected and subsequently isolated from engorged Stomoxys spp as well as from their proboscises. However, the question of LSDV viability in *Stomoxys* spp remains to be answered. Thus, the experiments reported here were designed to define the duration of LSDV retention in three *Stomoxys* species following intrathoracic inoculation.

## Materials and Methods

### Virus strain

In this experiment, LSDV isolates Kubash/KAZ/16 recovered from an infected cow during an outbreak of LSD in Atyrau, Kazakhstan in 2016 were used (9). The virus was grown on primary lamb testis (LT) and three times passaged in vitro using calves. Primary LT cell cultures were derived from pre-pubertal lambs according to Plowright and Ferris (10). Cell cultures exhibited 90% cytopathic effects (CPE), were freeze-thawed three times, and then centrifuged at 2000 g for 20 min to remove cell debris and stored at −70 °C until required.

### Insect colony

LSDV negative *Stomoxys* spp colonies used in this study were provided by the Entomology section of the Research Institute for Biological Safety Problems (RIBSP). They were maintained in the environmental chamber at 25°C and 80% humidity, and heparinized bovine blood soaked into cotton pads were offered twice daily at 9 a.m. and at 5 p.m.

### Design of experiment

As part of an ongoing study of the role of hematophagous flies in the transmission of LSDV in cattle in the South Kazakhstan, we utilized three laboratory reared *Stomoxys* species. The method of intrathoracic inoculation of LSDV in adult *Stomoxys* spp was conducted according to the protocol published by Rochon, Baker (11) and De Leon and Tabachnick (12). Prior to experiment commencement, approximately 200 five day old *Stomoxys* flies of each species (*Stomoxys Calsitrans, Stomoxys Sitiens, Stomoxys Indica)* were aspirated into separate plastic cages and then were temporarily anaesthetized using CO_2_ in a gas chamber and placed onto a light ice board and fixed using entomological forceps. Individual flies were inoculated in the dorsal area of the thorax in the junction of the scutum and the scutellum. Virus suspension of 6μl (Kubash/KAZ/16, 10^6^ TCID_50_) was administered using 100μl Bevel tip 1710N Series Hamilton Gastight microliter syringe (USA). A 26 gauge sterile removable needle was used and changed for each individual fly (Supelco Inc, Bellefonte, PA). Following virus inoculation flies were kept in separate cages and maintained at room temperature (24°C and 70% humidity).

*Stomoxys* flies were tested for the presence of LSDV nucleic acid by gel-based PCR and virus isolation (VI) at 0, 6h, 12h, 24h, 2d, 3d, 4d, 5d, 6d, 7d, 8d, 9d, 10d post intrathoracic inoculation (pii). Pools of ten flies each of each *Stomoxys* species inoculated as described above were collected and tested by PCR and VI immediately following virus administration (0hr). Prior to running tests, pools were rinsed with PBS solution to eliminate contamination. The remaining inoculated *Stomoxys* flies were placed in separate cages according to species and allowed to recover for 6 h. For sampling purposes, *Stomoxys* spp were aspirated from the cages with a vacuum, anaesthetized in a 5% CO_2_ gas chamber and transferred to a centrifuge tube and stored at −70°C until tested. Freeze–thawed flies were then washed three times in PBS and rinsed using distilled water to eliminate surface contamination, homogenized in 300μl Hanks solution, spun at 1500 g for 10 min and the supernatant was used for LSDV detection and isolation.

Additionally, three control groups of *Stomoxys* spp each containing 20 individuals of each species were inoculated by the method previously described to determine the total number of infected among inoculated flies as well as to determine the efficacy of the intrathoracic inoculation technique in use.

### Polymerase chain reaction

Virus amplification test was conducted utilizing protocol reported by Tuppurainen, Venter (13). For DNA extraction, a QIAamp DNA Kit (QIAGEN, USA) was used according to the manufacturer’s instructions.

For PCR assay, to produce 192 bp of amplified nucleotide reactions, the forward 5’-TCC-GAGCTC-TTT-CCT-GAT-TTT-TCT-TAC-TAT-3’ and reverse 5’-TAT-GGT-ACC-TAA-ATT-ATA-TACGTA-AAT-AAC-3’ primers were utilized (14). The parameters for DNA amplification in a Thermal Cycler (Eppendorf Mastercycler) were as follows: 95 °C for 2 min, 95 °C for 45 s, 50 °C for 50 s, 72 °C for 1 min (34 cycles), and 72 °C for 2 min. The PCR products obtained were loaded in 1.5% agarose gel electrophoresis in the presence of *Ethidium bromide*, and the results visualized using Bio-Imaging Systems MiniBIS Pro (Israel).

### Virus Isolation

Virus isolation was performed according to standard operational procedures (SOP) of the BSL-3 laboratory of the RIBSP, based on OIE (15) manuals. Briefly, 1 mL supernatant were inoculated on to lamb testes cells in 25 cm^2^ cell culture flasks and allowed to incubate at 37 °C for 1 h. Following incubation, culture media was rinsed with PBS and overlaid with Glasgow’s Minimal Essential Medium containing 0.1% penicillin, 0.2% gentamycin and 2% foetal calf serum. The cell monolayer was monitored daily for characteristic CPE. In the case no CPE was observed, the cell culture was freeze–thawed three times and the two or three blind passages were conducted. The culture media used were kept at −70 °C until needed. Cell culture flasks exhibiting CPE were tested with gel-based PCR to confirm that CPE was caused by LSDV.

## Results

### Demonstration of LSDV retention in Stomoxys Calcitrans

Virus retention following intrathoracic inoculation procedures resulted in infection in 95% of inoculated flies (Table 1). Pools samples collected for PCR were tested positive at time intervals 0, 6h, 12h, 1d, 2d, 3d, 4d, 5d, 6d and 7 days post intrathoracic inoculation (pii) whereas samples tested for virus isolation were LSDV positive with characteristic CPE from time interval 0 up to 24h pii (Table 2).

**Table 1.** Summary obtained from individual samples of Stomoxys species.

| Stomoxys<br>species | Cohort<br>size/individuals<br>inoculated | PCR<br>positive | % infected |
| --- | --- | --- | --- |
| *Stomoxys*<br>*Calcitrans* | 20 | 19 | 95 |
| *Stomoxys*<br>*sitiens* | 20 | 18 | 90 |
| *Stomoxys*<br>*indica* | 20 | 19 | 95 |

**Table 2.** PCR and virus isolation results of *Stomoxys spp* at different time intervals following intrathoracic inoculation with LSDV.

| Time/<br>day<br>Post<br>inocul<br>ation | <u>*Stomoxys Calcitrans*</u> |  |  | <u>*Stomoxys Sitiens*</u> |  |  | <u>*Stomoxys Indica*</u> |  |  |
| --- | --- | --- | --- | --- | --- | --- | --- | --- | --- |
|  | PCR(no<br>positive<br>/no<br>tested)* | Virus<br>isolation<br>(no<br>positive/no<br>tested)* | Virus<br>isolation<br>(TCID<sub>50</sub>/mL), mean | PCR(no<br>positive<br>/no<br>tested)* | Virus<br>isolation<br>(no<br>positive/n<br>o tested)* | Virus<br>isolation<br>(TCID<sub>50</sub>/mL), mean | PCR(no<br>positive/n<br>o tested)* | Virus<br>isolation<br>(no<br>positive/no<br>tested)* | Virus<br>isolation<br>(TCID<sub>50</sub>/mL), mean |
| 0h | 10/10 | 7/10 | 4.5 | 9/10 | 9/10 | 4.1 | 10/10 | 7/10 | 3.9 |
| 6h | 8/10 | 4/10 | 3.2 | 9/10 | 5/10 | 2.8 | 8/10 | 6/10 | 2.5 |
| 12h | 7/10 | 2/10 | 2.0 | 8/10 | 4/10 | 1.3 | 6/10 | 3/10 | 1.3 |
| 1 | 7/10 | 2/10 | 1.1 | 6/10 | 1**/10 | - | 5/10 | 1**/10 | - |
| 2 | 7/10 | 1/10 | - | 6/10 | 0/10 | - | 5/10 | 0/10 | - |
| 3 | 6/10 | 0/10 | - | 5/10 | 0/10 | - | 4/10 | 0/10 | - |
| 4 | 5/10 | 0/10 | - | 4/10 | 0/10 | - | 3/10 | 0/10 | - |
| 5 | 3/10 | 0/10 | - | 3/10 | 0/10 | - | 3/10 | 0/10 | - |
| 6 | 1/10 | 0/10 | - | 1/10 | 0/10 | - | 2/10 | 0/10 | - |
| 7 | 1/10 | 0/10 | - | 0/10 | 0/10 | - | 1/10 | 0/10 | - |
| 8 | 0/10 | 0/10 | - | 0/10 | 0/10 | - | 0/10 | 0/10 | - |
| 9 | 0/10 | 0/10 | - | 0/10 | 0/10 | - | 0/10 | 0/10 | - |
| 10 | 0/10 | 0/10 | - | 0/10 | 0/10 | - | 0/10 | 0/10 | - |

\*- number of positive and number of tested pools
\*\*- CPE after second blind passage in (LT) cell culture

### Demonstration of LSDV retention in Stomoxys Sitiens

LSDV DNA retention was detected in 90% of individually infected flies (Table 1). PCR assay was positive for the presence of LSDV in pools samples taken in time intervals 0, 6h, 12h, 1d, 2d, 3d, 4d, 5d and 6 days pii (Table 2). Virus isolation was carried out in cell culture, and CPE characteristic to LSDV was observed in time intervals 0, 6h, 12h pii. Additionally, following the second blind passage virus was recovered from pool samples collected on day 1 pii (Table 2).

### Demonstration of LSDV retention in Stomoxys Indica

At this stage, an infection rate of 95% was demonstrated among intrathoracically infected individuals (Table 1). Viral nucleic acid was detected in time intervals 0, 6h, 12h, 1d, 2d, 3d, 4d, 5d, 6d and 7 days pii (Table 2). CPE similar to LSDV was seen in time intervals 0, 6h, 12h pii while on day 1 pii virus was isolated following the second blind passage in the (LT) cell culture (Table 2).

## Discussion

In this study, the intrathoracic injection of three *Stomoxys* species was demonstrated to establish the viability of LSDV in the haemolymph of flies. When LSDV was administered into the hemocoel of adult *Stomoxys* flies, the virus titre decreased over time (Table 2), which contrasts with the findings reported by Rochon, Baker (11). In this study, the replication of LSDV was not observed in insect tissue which is in agreement with reports published by Chihota, Renniet (16) and Issimov, Kutumbetov (7). However, the haemolymph does not appear to be inimical to the virus as the alimentary track as demonstrated by relatively prolonged viral DNA presence in intrathoracically inoculated flies. This observation was similar to those described by Schurrer, Dee (17) and Rochon, Baker (11). All three *Stomoxys* species were capable of virus retention after intrathoracic inoculation either in groups (pools) or in individuals. Moreover, viable LSDV was recovered from *Stomoxys spp* up to 48h following virus injection, whereas viral nucleic acid was detected up to day 7 post intrathoracic inoculation.

In a previous study of Porcine reproductive and respiratory syndrome virus retention in house flies, the factors found to influence pathogen stability following *per os* acquisition have been explored. Thus, time and ambient environment have a detrimental effect resulting in virus concentration decrease over time in the midgut environment (17).

In blood-feeding insects, the midgut barrier is a significant line of defense against infection as most of the pathogens are ingested orally via contaminated bloodmeal (18). The midgut epithelial cells are refractory to infection, and their immunity to viruses is genetically controlled (19). In addition to this, nonimmune and refractory *individuals* can arise within the same cluster of insects (20). Prior research has indicated that previously refractory insects may become competent vectors, in the event that a pathogen overcomes the midgut barrier (12). Studies on the role of *Culicoides* in the transmission of bluetongue virus (BTV) reviled that a colony of intrathoracically BTV infected midges were able to transmit the virus in their saliva. This, in turn, indicates a breach of salivary gland barrier and its susceptibility to infection (20). Similarly, the integrity of the midgut barrier can also be affected by microfilariae infection and high doses of B. thuringiensis (21, 22). These findings demonstrate the likelihood that some individuals of insect populations might become permissive enough to allow a new dynamic of the vector – pathogen system (23). In conclusion, the outcomes obtained in this study illustrate the incompetence of *Stomoxys* species to serve as biological vectors of LSDV however further experiments with intrathoracically inoculated flies are of importance to determine if there is a salivary gland barrier for LSDV in the *Stomoxys* fly.

## Acknowledgements

This study was funded by “Bolashak” international scholarship program as well as by research project # AP05135323 of the Science Committee/ Ministry of Education and Science/Republic of Kazakhstan.

## Conflict of interest

The author declare that they have no conflict of interest.

## Data availability

The data obtained during this study are openly available in KNB Data Repository at doi:10.5063/F1PN9408.

